# Spatial patterns in the diet of *Gyps* vultures in India and their implications for conservation

**DOI:** 10.1101/2023.08.21.554114

**Authors:** Mousumi Ghosh-Harihar, Nidhi Yadav, Nehal Gurung, Darshan C.S., Shashikumar B., Vishnudas C.K., Vibhu Prakash, Rhys E. Green, Uma Ramakrishnan

**Author notes:** Corresponding author: Mousumi Ghosh-Harihar, National Centre for Biological Sciences, Tata Institute of Fundamental Research, Bellary Road, Bengaluru-560065, Karnataka, India. Nature Conservation Foundation, “Amritha”, 1311, 12^th^ Cross, Vijaynagara, 1^st^ Stage, Mysore-570017, India (Mousumi Ghosh-Harihar). (MGH), (UR), (NY), (NG). (DCS); (SB).

## Abstract

Dietary analyses are particularly useful for developing conservation programmes for species threatened by resource depletion, poisoning and environmental pollution. *Gyps* vultures in South Asia represent one such case, having undergone a population collapse caused by feeding on carcasses of cattle treated with the non-steroidal anti-inflammatory drug diclofenac. Following a ban on the veterinary use of diclofenac, populations of vultures remain low and mostly concentrated near protected areas. Understanding the role of protected areas in the recovery of these critically endangered species requires analyses of spatial variation in their diet. We used faecal metabarcoding to investigate the spatial variation in the diet of *Gyps* vultures across four landscapes from sites located inside and outside protected areas. We collected faecal 642 samples, of which 419 yielded adequate molecular data to identify the vulture species and 30 molecular operational taxonomic units corresponding to at least 28 diet species. The diet was dominated by large ungulates and varied across landscapes, protection status and vulture species, with the observed variation largely explained by differential intake of wild and domestic species. Domestic livestock was present in >95% of the samples from central, west and north India, but 77% of the samples had only wild species in south India. This variation was explained by livestock density within 100 km radius of the sampling site. Our results imply that protected areas may not offer a respite from possibility of diclofenac-poisoning across most parts of the country and efforts should continue unabated to remove nephrotoxic drugs from veterinary use.

## 1. Introduction

Reliable data on dietary habits of species provide key insights into their evolutionary adaptations as well as ecological and trophic interactions with their environment (Gainsbury et al., 2018; Kissling et al., 2014; Symondson, 2002). Dietary analyses can be particularly useful in identifying targeted conservation actions for endangered species whose decline is caused by depletion of prey/resources, poisoning and effects of toxic environmental pollutants (Liu et al., 2018; Oehm et al., 2011; Sullins et al., 2018). The catastrophic decline of Old World vultures in the Indian subcontinent is such a case, in which a severe crash in populations was tightly linked to their diet (Oaks et al.; 2004; Green et al., 2004; Shultz et al., 2004). Global populations of three species in the genus *Gyps* endemic to South and Southeast Asia (White-rumped Vulture *G. bengalensis*, Indian Vulture *G. indicus* and Slender-billed Vulture *G. tenuirostris*) declined so rapidly that all are now listed as Critically Endangered on the IUCN Red List (Birdlife International 2023).

Up to the mid-1990s, carcasses from the large population of domesticated cattle (*Bos indicus, B. taurus, B. indicus* x *taurus* hybrids and *Bubalus bubalis*) in India supported at least millions of the three resident *Gyps* species (Prakash et al. 2019), with White-rumped Vulture thought to be the most abundant large bird of prey in the world (Houston, 1985). Due to religious and cultural limits on slaughter for human consumption, vast amounts of cattle carrion were available to the vultures for consumption and probably constituted the principal and most predictable food source (Cuthbert et al., 2014). However, beginning in about 1994 (Cuthbert et al. 2014), the region’s three resident *Gyps* vultures suddenly underwent catastrophic population declines (99.9% for White-rumped Vulture), necessitating urgent species recovery measures (Prakash et al., 2003). These declines prompted the IUCN-World Conservation Union to list the three species as Critically Endangered in 2000 (Hilton-Taylor, 2000). The cause was identified to be the veterinary use of the non-steroidal anti-inflammatory drug (NSAID) diclofenac to provide palliative care to dying cattle using on multiple lines of evidence (Oaks et al., 2004; Green et al., 2004; Shultz et al., 2004). Birds developed visceral gout associated with kidney failure and died within days when they fed on domesticated ungulates containing diclofenac residues. The *Gyps* vultures are particularly prone to extinction due to their long generation lengths, larger body sizes, preference for large mammal carcasses that are more likely to be accidentally or deliberately contaminated with poisons, and social foraging habits, which make a single poisoned carcass adequate to kill many (even hundreds) of birds (Buechley and Şekercioğlu, 2016). In fact, it is estimated, based upon population modelling, that just 0.8% of the available cattle carcasses would need to have been contaminated with lethal levels of diclofenac to fully explain the rapid rate of decline of the White-rumped Vulture (50% decline per year) observed in both India and Pakistan (Green et al., 2004). When the proportions of cattle carcasses in India contaminated with diclofenac were measured directly by taking liver samples (Green et al. 2007) it was found that contamination levels were more than sufficient to fully account for the observed decline.

Recognizing this threat, veterinary use of diclofenac was banned in India, Nepal and Pakistan in 2006 and in Bangladesh in 2010. Although the proportion of contaminated carcasses was reduced by half in India within four years (Cuthbert et al., 2014), illegal veterinary use of diclofenac has continued in India. Undercover surveys of veterinary pharmacies showed that diclofenac is still offered for sale for the treatment of cattle at many pharmacies, with availability showing very variable trends over time in different Indian states (Galligan et al. 2021). *Gyps* populations in India have been approximately stable since the late 2000s (Prakash et al. 2019), but numbers remain low, with no clear signs of recovery. In Nepal, where veterinary use of diclofenac has now been almost eliminated by awareness-raising campaigns, *Gyps* populations have been increasing at the maximum possible rate since about 2012 (Galligan et al. 2020). Road transect surveys in India showed that *Gyps* vulture had declined to a smaller extent over the same time period near to some National Parks (protected areas) than in other areas (Prakash et al 2012). In addition, road transect surveys in India during 2003-2015 showed that the vultures remaining after the decline were much more abundant within about 100 km of National Parks than further away (Prakash et al., 2012). It was hypothesized that National Parks might offer more uncontaminated wild ungulate carcasses and thereby, reduce exposure to NSAIDs and perhaps also reprisal poisoning. However, confirming this hypothesis requires a reliable dietary data from wild vultures both inside and outside these protected areas. Furthermore, the observed diet needs to be related to relevant ecological covariates including relative availability of domestic livestock and wild ungulate species. This is particularly crucial since wild prey densities vary considerably among different protected areas within India owing to differences in environmental factors and management. Many protected areas also face substantial grazing pressure due to domestic/feral livestock ranging from surrounding villages. The number of cattle grazing within and around National Parks could be as high as 200,000 as seen in the case of Sariska National Park in north-western India (Bindra, 2017). Presence of domesticated ungulates in the diet continues to present a significant threat to wild vultures in India owing to weak enforcement and continued illegal use of diclofenac and legal veterinary use of other nephrotoxic NSAIDs (e.g. ketoprofen, nimesulide and aceclofenac) (Cuthbert et al., 2016; Galligan et al., 2021). Therefore, a better understanding of the spatial variation in vulture diet and its ecological determinants, particularly with reference to availability of cattle carcasses, will be particularly critical for prioritizing *in situ* conservation actions, identifying relatively safe areas for reintroduction and ensuring persistence of the gradually recovering vultures.

Vulture diet has been evaluated previously by examination of undigested material (hair, bones, scales etc.) in regurgitated pellets (Donázar et al., 2010) or food remains under nests (Blanco et al., 2019). However, this method may underestimate domestic carcasses in areas where skinners remove the hides of cattle before they are made available to scavengers (a practice common in many parts of the subcontinent), due to less identifiable hair being present (Donázar et al., 2010; Kelly et al., 2007). DNA-based diet analysis presents an alternative to overcome this limitation, and can be more efficient, detect more prey items and distinguish better between prey species (Casper et al., 2007). In a previous study, we developed a standardized diet metabarcoding pipeline to simultaneously identify the *Gyps* species, its sex and dietary species, which makes the diet analyses both efficient and more reliable (Ghosh-Harihar et al., 2021).

In the present study, we generated dietary data from wild *Gyps* vultures across multiple sites in India, both inside and outside protected areas and explored the ecological correlates of the diet, particularly with reference to presence of livestock. We characterized the dietary habits of wild vultures from 642 faecal samples collected from nesting/roost sites and analyzed each faecal sample using DNA metabarcoding and high throughput sequencing. Using a complement of four primer sets, we were able to perform parallel identification of the *Gyps* species, sex and the species represented in the diet. We then tested for spatial variation in diet composition of these species and examined the ecological predictors explaining the pattern. Finally, to ascertain if protected areas influenced intake of domesticated ungulates carcasses, which might be tainted with toxic drugs, we built models to identify the factors that best explained the observed spatial variation in probability of domesticated ungulates being present in diet of wild *Gyps* species.

## 2. Materials and methods

### 2.1 Study area and sample collection

We sampled vulture faeces across multiple locations, both inside and outside protected areas, in the states of Himachal Pradesh (North India), Rajasthan (Western India), Madhya Pradesh (Central India), Kerala, Karnataka and Tamil Nadu (South India) during May 2018-January 2022 (Fig. 1, Table A1). These locations were selected based on records of recent breeding (last five years) using eBird records and personal communication with local conservationists and naturalists. We visited these locations during the non-breeding season (November-April) and located active nesting and roosting sites of White-rumped Vultures (*Gyps bengalensis*) and Indian (Long-billed) Vultures (*G. indicus*) with help from local forest officials and trackers. We found White-rumped vultures nesting on tall trees, while Indian vulture nests were found on steep cliff faces. In addition, we found both species roosting together on trees in some locations, often with migratory griffons (Eurasian-*G. fulvus* and Himalayan-*G. himalayensis*).

**Figure 1.**
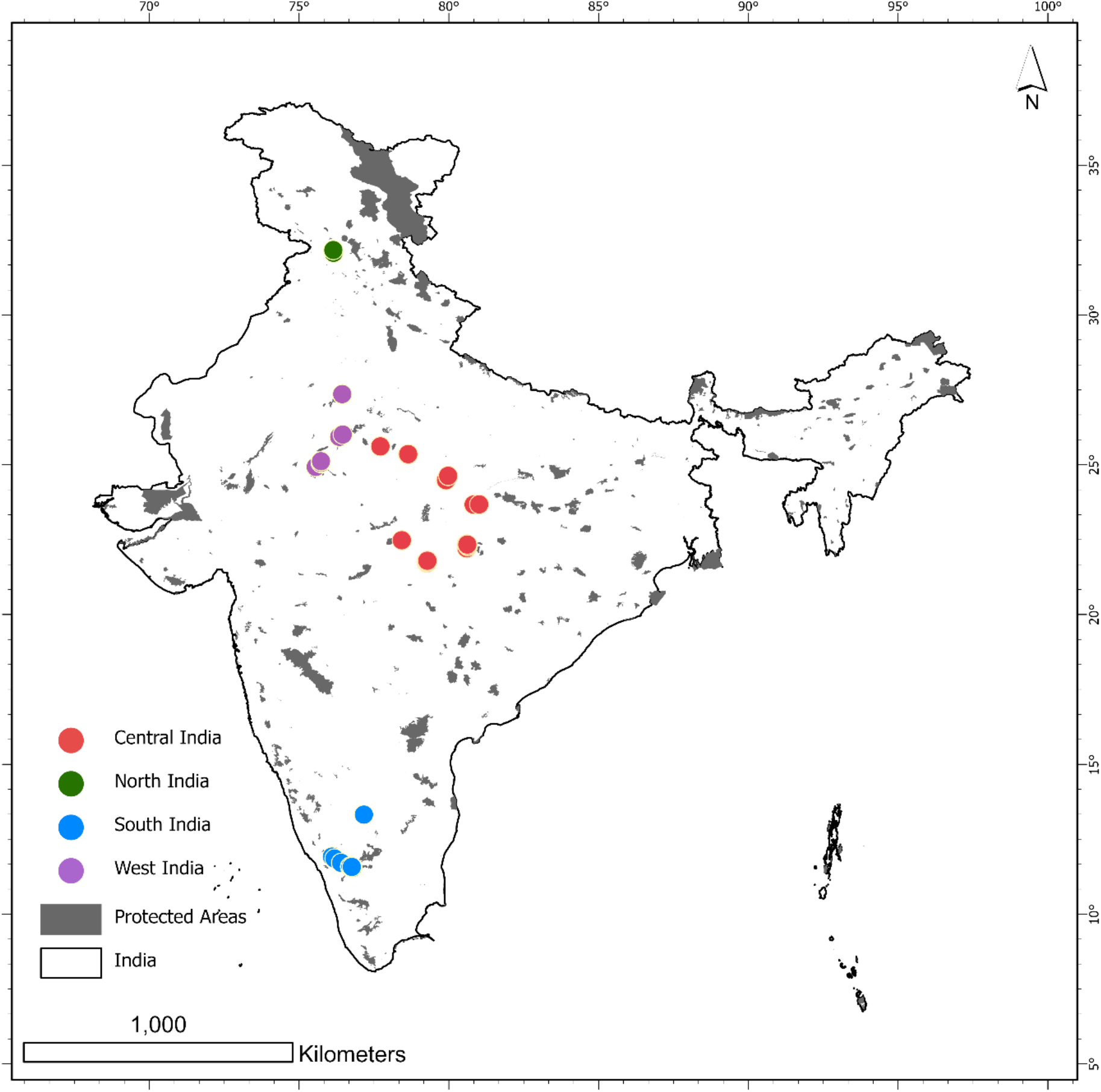
The sampling locations for collection of vulture droppings (*N = 642)* across four landscapes both inside and outside protected areas (shaded in grey) in India.

Once a roost/nest of the target species was located, we collected relatively fresh faecal samples (determined visibly) from various substrates (soil, leaves, tree bark, rocks) directly under the nest/roost. We collected samples that were spatially separated to minimize chances of sampling the same individual repeatedly and increased the chance that the sample came from one individual by avoiding overlapping droppings. Vulture droppings are composed of both urine (principally uric acid, which is white) and faecal material (which is dark). The dark portion was swabbed using a sterile plastic applicatory rayon swab dipped in Longmire lysis buffer (Longmire et al., 1997). We broke the tip and collected it in a 2 ml vial containing 1 ml of the same buffer solution. Sterile disposable gloves were used while collecting faecal samples to minimize the risk of contamination with human DNA. Similarly, we collected field negative controls, where a nearby substrate (without droppings) was swabbed. Each sample was labeled with a unique identification code. Collected samples were stored in the field at room temperature and transferred to the laboratory at National Centre for Biological Sciences at Bangalore within a month and stored at -20°C until DNA extraction.

### 2.2 Dietary data generation

The detailed methods for the diet data generation are described in Ghosh-Harihar et al. (2021). We extracted faecal DNA from the field samples following the manufacturer’s recommendations using the QIAmp Fast DNA Stool Mini Kit (QIAGEN, Germany). Following extraction, we amplified each sample using the *CytbV1* primer and visualized the amplification using gel electrophoresis on 2% agarose gel to confirm the presence of *Gyps*-specific DNA. Thereafter, we amplified the *Gyps-*positive samples using two separate multiplex PCR reactions to allow parallel identification of vulture species (*CytbV1*), vulture sex (*VsexZW*) and prey species (*12sV5* with the blocker *GypsB; 12SV-mod*) (Table A2). All primers were modified by adding the Illumina overhang adapters. We amplified the samples, including several negative (field, extraction, PCR) and positive (mock community sample) controls. Final libraries (including indexing controls) were sequenced with an Illumina MiSeq (Illumina, USA). We analyzed the HTS data separately for each of the fours metabarcodes following Ghosh-Harihar et al. (2021).

For each metabarcode, we first collapsed the final set of variants into discrete unique molecular taxonomic units (MOTU) using a 2% sequence divergence threshold and relative abundance of the sequences. We resolved the taxonomy of variants with >2% divergence with the help of the topology of the trees generated from additional phylogenetic analyses. To confirm species-level identity, we performed additional BLAST hits in GENBANK with very stringent criteria (E-value threshold: 1e− 10; minimum query coverage: 98%). Biogeographic information on species occurrence in the region was also used to ascertain diet species identify in some cases. The metabarcodes could not differentiate between the different species of domestic cow (*Bos indicus, B. taurus, B. indicus* x *taurus* hybrids) all of which were assigned as *B. taurus*, and we used the assignment as such for our purposes. However, differentiating between domestic cow and gaur (*B. gaurus*), which is a wild ungulate and represents safe food source for the vultures, was critical. To differentiate between domestic cow and wild gaur (*Bos gaurus*), the positive controls contained DNA from *Bos gaurus*. The unique *Bos* sequence variant from the positive control was then used as the reference to identify *B. gaurus* sequences in the samples collected from the wild (Appendix A). We excluded sequences identified as bats, swifts, human and rodents as contaminants based on sequences obtained from field negative controls. Since sex could not be determined for a considerable number of samples from one specific site, the influence of sex on diet was not investigated.

### 2.3 Environmental variables

We considered multiple environmental variables, which could potentially influence the diet, capturing spatial variation for a 100-km radius around the study site coordinates. This radius was chosen because vulture abundance is highest within 100 km of National Parks (Prakash et al. 2012) and because the primary *in situ* conservation strategy in South Asia for recovering and conserving remaining wild populations of these species involves securing Vulture Safe Zones (VSZ) with a radius of 100 km centered around known breeding sites (Mukherjee et al., 2014). Concerted efforts are being made within these VSZs to remove toxic NSAIDs from the vulture food supply through targeted persuasion, education, and awareness programmes to allow recovery and create ‘safe’ reintroduction sites. We considered multiple variables including summed livestock (cows, buffaloes, goat, sheep) density extracted from the Gridded Livestock of the World (GLW v2.01; Gilbert et al., 2018; livestock.geo-wiki.org), mean annual precipitation, mean annual temperature (worldclim.org), mean elevation, mean terrain ruggedness index (TRI; Wilson et al., 2007), tree cover (Global Land Cover Database, 2003), area under protected area coverage (Ghosh-Harihar et al., 2019) and mean human impact index (HFI, Venter et al., 2016). For these continuous variables, we calculated the sum or mean of all pixels partially or wholly within the 100 km radius and mean-standardized the values, so that coefficients were directly comparable as effect sizes. To minimize collinearity, we began with a full model including all covariates and sequentially dropped the covariates with the highest variance inflation factor (VIF). We recalculated the VIFs and repeated this process until all VIFs were smaller than a pre-selected threshold of 2 to enable detection of even weaker ecological signals (Zuur et al. 2010).

### 2.4 Diet composition analyses

We applied non-metric multidimensional scaling (nMDS) ordination to visualize the variation of diet composition among sites distributed across the four landscapes (North India, West India, Central India and South India) from samples collected from within and outside protected areas (Fig. 1). For this study, nMDS was performed whereby the ordination was constructed from the Jaccard dissimilarity matrix of pairwise dissimilarities between samples based on presence/absence data. The nMDS was performed using the function “metaMDS” from *R* package *vegan* (Oksanen et al., 2019). The nMDS ordination projects the multivariate data into a space with a smaller number of axes in such a way that the most similar samples are close together, while samples that are more different are further apart. We fitted linear vectors into the nMDS ordination to examine the relationship between environmental predictors and the diet composition using the function *envfit* in the *vegan* package of *R*. This was done by running first an NMDS ordination of diet species and then followed by fitting the linear vectors (environmental variables) simultaneously. The analysis used the same species diet species presence/absence data. The significance of the environmental vectors was evaluated using a permutation test with 1,000 random permutations. To test for differences in diet composition across the landscapes and between samples collected from within protected areas and outside and among species, we performed the Analysis of Similarities (ANOSIM) permutation tests in *vegan* package of *R* (Oksanen et al., 2019), with 999 random permutations based on the Jaccard dissimilarity matrix (with Bonferroni corrected p-values). Finally, a similarity percentage analysis (SIMPER) was used to assess which diet species contributed most to the observed dissimilarity between these groups.

### 2.5 Ecological correlates of domesticated ungulates in the diet

To visualize the links between livestock in diet, the landscape and the site being inside or outside protected area, we constructed Sankey diagram with the help of the “*ggalluvial*” and “*ggplot2*” packages in R. We analyzed the environmental variables that influenced the presence of domestic ungulates (cow, buffalo, camel, goat and sheep) in the diet of *Gyps* species (a binary variable) using generalized linear mixed models (GLMM) with a binomial error distribution and logit link function. To avoid faecal samples from the same individual, we sampled in different areas (at intervals of more than 200 m), and only one sample was collected at the same site and time. For samples collected within 500m of each other, we assigned a common site ID (sites=30), which was used as a random effect in these models to account for repeated measurements. Vulture species identity was also included as a categorical fixed effect. After testing for multicollinearity among the variables, we retained summed livestock density, area under protected area coverage, mean human impact index and mean annual precipitation as explanatory variables in our models. We also included a two-way interaction between summed livestock density and area under protected area coverage. All the candidate models were called in the *lme4* package (Bates et al., 2015) in R (version 4.1.3). We ranked the models based on size-corrected AIC values and model with ΔAIC_c_ of <2 from the best fit model were considered statistically indistinguishable (Anderson and Burnham, 2002). Environmental variables were considered supported if their coefficient CI did not span zero. We detected no spatial autocorrelation among the residuals using Moran’s I test (*p* > 0.05).

## 3. Results

### 3.1 Sampling overview

We collected 642 faecal samples targeting the *Gyps* species from nesting and roosting locations depicted in figure 1. Of these, 425 samples were found to be vulture-positive and processed further to identify the species, sex and diet. After applying all the quality filters, three samples did not have adequate number of sequences to determine species identity, a single sample came from Red-headed vulture (*Sarcogyps calvus*), and no diet species could be detected in two other samples. These samples were excluded from further analysis. The remaining 419 samples were identified as belonging to Indian vulture (*N*= 241), White-rumped vulture (*N*= 142), Eurasian griffon (*N*=22) and Himalayan griffon (*N* = 14). Our study did not locate any samples from Slender-billed vultures, which are principally found in West Bengal and Assam.

### 3.2 Dietary composition

We detected 30 distinct dietary MOTUs in the faecal samples belonging to four *Gyps* species using DNA metabarcoding (Table A1). Of these, we could assign species level identification to 26, two could be identified to the genus level (*Urva sp*.; *Prionailurus sp*.) and two MOTUs could be resolved only up to subfamily level (Cervinae, Felinae). Since the metabarcodes could not differentiate between wild pigs and domestic pigs, we did not include them in domestic livestock for the purposes of this study. Apart from mammalian species, we also detected Indian peafowl, red jungle fowl and grey jungle fowl in the diet of these species.

The mean number of diet MOTUs per sample was 2.86 ± 1.59 *SD* (range 1–9). The five most commonly detected MOTUs were domestic cow (*Bos taurus*, 79%), domestic buffalo (*Bubalus bubalis*, 47.9%), chital (*Axis axis*, 37.8%), Cervinae (16.7%), sambar (*Rusa unicolor*, 15.7%) and nilgai (*Boselaphus tragocamelus*, 15.7%). Those rarely encountered included 23 species, which were detected in less than 10% of the samples (Table A1). The diet of the two resident species with larger sample sizes represented almost all the dietary MOTUs (Indian vultures-26; White-rumped vulture-25), while the other two migratory species with fewer samples also showed a smaller, though considerable, number of dietary MOTUs (Himalayan Griffon-13; Eurasian Griffon-18).

The nMDS ordination plot showed that samples collected in South Indian sites showed a diet composition distinct from the samples from other sites (Fig. 2). Of the four environmental variables, three were significantly correlated with the nMDS axes: summed livestock density (*envfit r*^*2*^=0.38, *P*=0.001) and area under protected area coverage (*envfit r*^*2*^=0.25, *P*=0.001) and mean terrain ruggedness index (*envfit r*^*2*^=0.08, *P*=0.001). Diet composition differed significantly across landscapes (ANOSIM *R= 0*.*3, P=001)*, between sites located within and outside protected areas (ANOSIM *R= 0*.*09, P=001)* and among species (ANOSIM *R= 0*.*21, P=001*). The SIMPER analysis indicated that domestic cow, water buffalo and chital were dominant contributors to observed dissimilarities between landscapes, species as well as inside and outside protected areas (Fig. 3, Table A3), along with other commonly detected species. For instance, a higher proportion of samples from South Indian samples contained wild ungulates and very few contained domestic species, whereas other landscapes had between 87-100% of the samples containing domestic cow and between 40%-61% showed the presence of water buffaloes (Fig. 3). Overall, samples from within protected areas contained more wild ungulates (e.g. chital-53% inside and 15% outside) and fewer domestic species; among the *Gyps* species, White-rumped vultures showed a lower proportion of samples with domestic cow (52.3%) and water buffalo (30%) than the other vulture species (domestic cow-91-93%).

**Figure 2.**
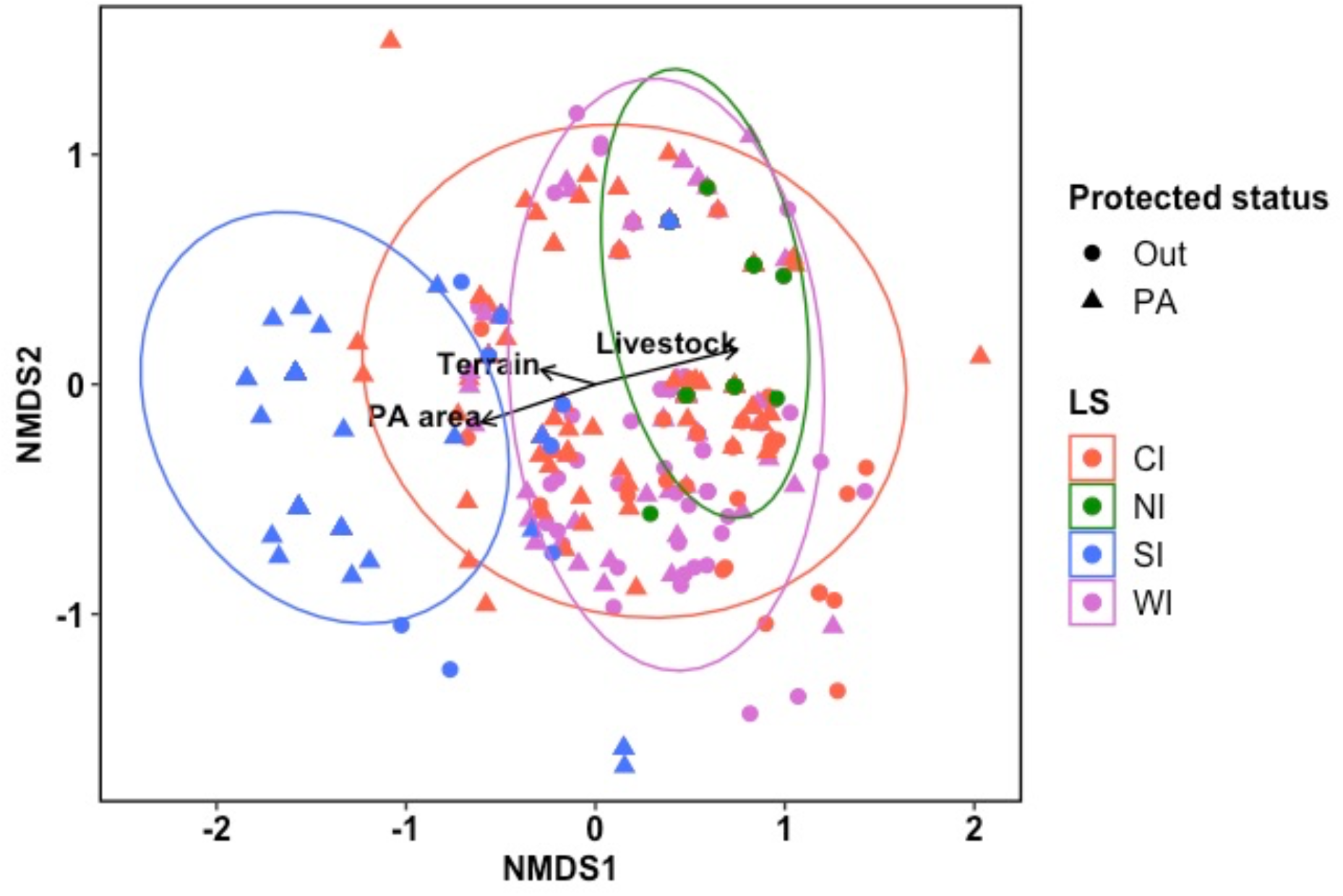
Non-metric multi-dimensional scaling (nMDS) plot showing the spatial variation in diet composition for *Gyps* vultures (*N* = 419) across four landscapes, and across protected areas (PA) and outside (Out) in India. 2D Stress value 0.135. Label shapes indicate if the sampling site is within or outside protected area, colour-coded by landscape. The vectors represent the environmental variables with significant Pearson correlation with the nMDS axes..

**Figure 3.**
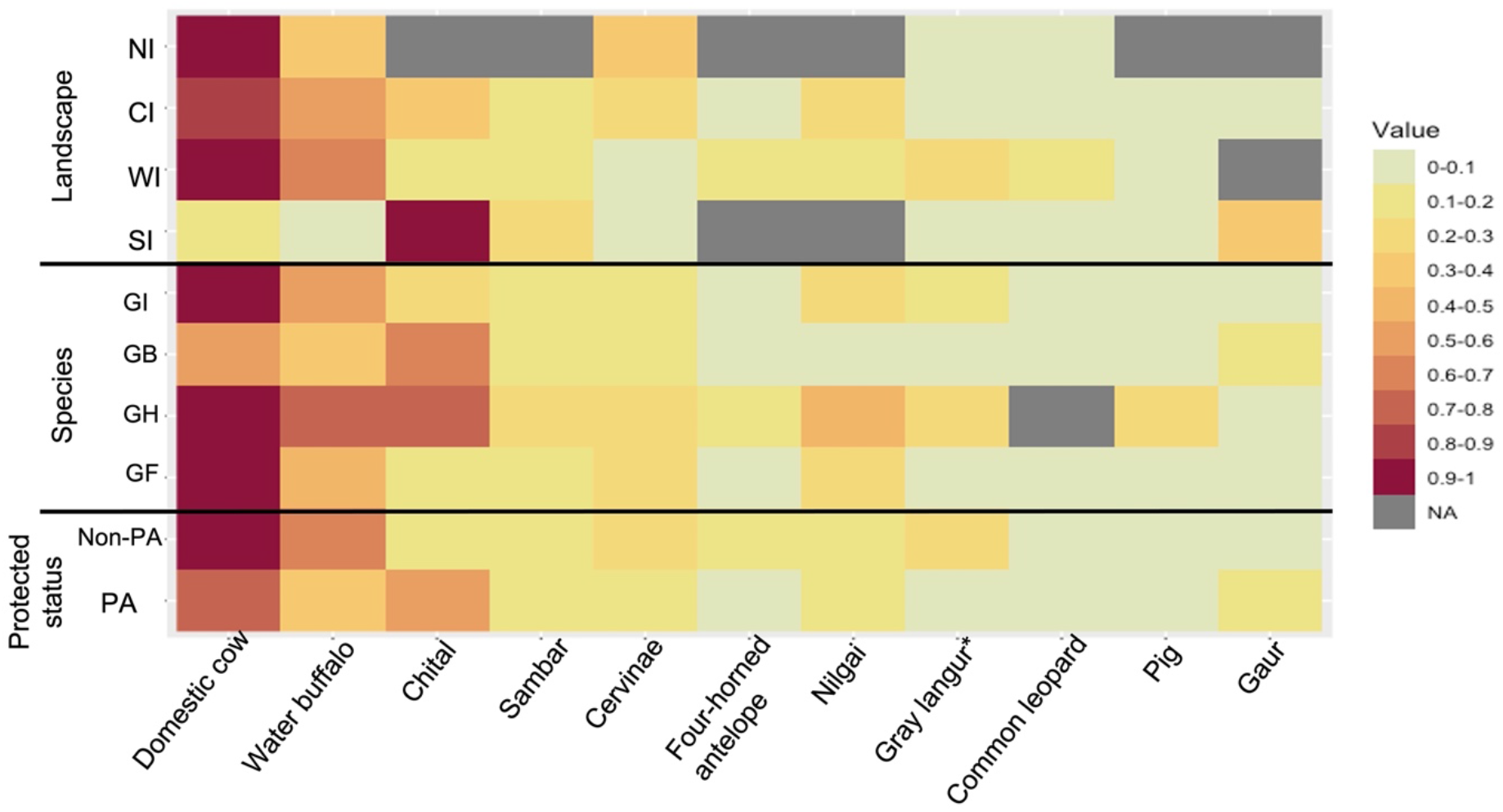
Heatmap representing the relative proportion of diet MOTUs that contributed to the differentiation of groups (SIMPER analysis): (a) landscapes, (b) species and (c) between sites situated within and outside protected areas. Only MOTUs that together contributed to > 70% of the differentiation are included.

### 3.3 Domesticated ungulates in diet

Only 23% of the samples from South India contained domesticated ungulates compared to the other three landscapes, where almost every sample had domesticated ungulate presence (Central India-95%, West India-99% and North India-100%) (Fig. 4). Across all landscapes, the samples with only wild species in diet came from sites located within protected areas. To identify the variables influencing the domesticated ungulate presence in the diet of *Gyps* species, we examined the parameter estimates from the GLMMs (Table A4). The best model (lowest AICc) included the additive effects of summed livestock density and area under protected area and an interaction between these two (Table 1). Summed livestock density had a positive influence on presence of domestic ungulates in the diet of vultures. The positive interaction term further implied that particularly in sites with high summed livestock density, area under protected area provided no respite from possibility of feeding on potentially poisoned domestic ungulate carcasses. The next best model (within 2 !AICc) also included human footprint index as an additional covariate, but the confidence interval of the estimate spanned zero.

**Table 1.** Parameter estimates of the top (with the lowest ΔAIC_c_) generalized linear mixed effect model. The GLMM for presence of domestic ungulates in diet of *Gyps* vultures was fitted with a binomial distribution and a logit link function, with site ID as a random effect (*N=*419 from 30 sites). Summed livestock density and the area under protected area were measured from within a radius of 100 km^2^ centred at each sampling site and mean standardized. Significant model parameters are highlighted in bold. The complete model set is presented in Table A4.

| | Estimate | SE | z-value | $P(z)$ |
| --- | --- | --- | --- | --- |
| Fixed effects |  |  |  |  |
| Intercept | 6.61 | 1.63 | 4.04 | <0.0001 |
| **Summed livestock density** | **4.45** | **1.39** | **3.2** | **0.001** |
| Area under protected area | 2.25 | 1.53 | 1.47 | 0.142 |
| **Summed livestock density x Area under protected area** | **2.37** | **0.97** | **2.43** | **0.015** |
| Random effects |  |  |  |  |
|  | Variance | SD |  |  |
| Site ID (Intercept) | 5.74 | 2.4 |  |  |

**Figure 4.**
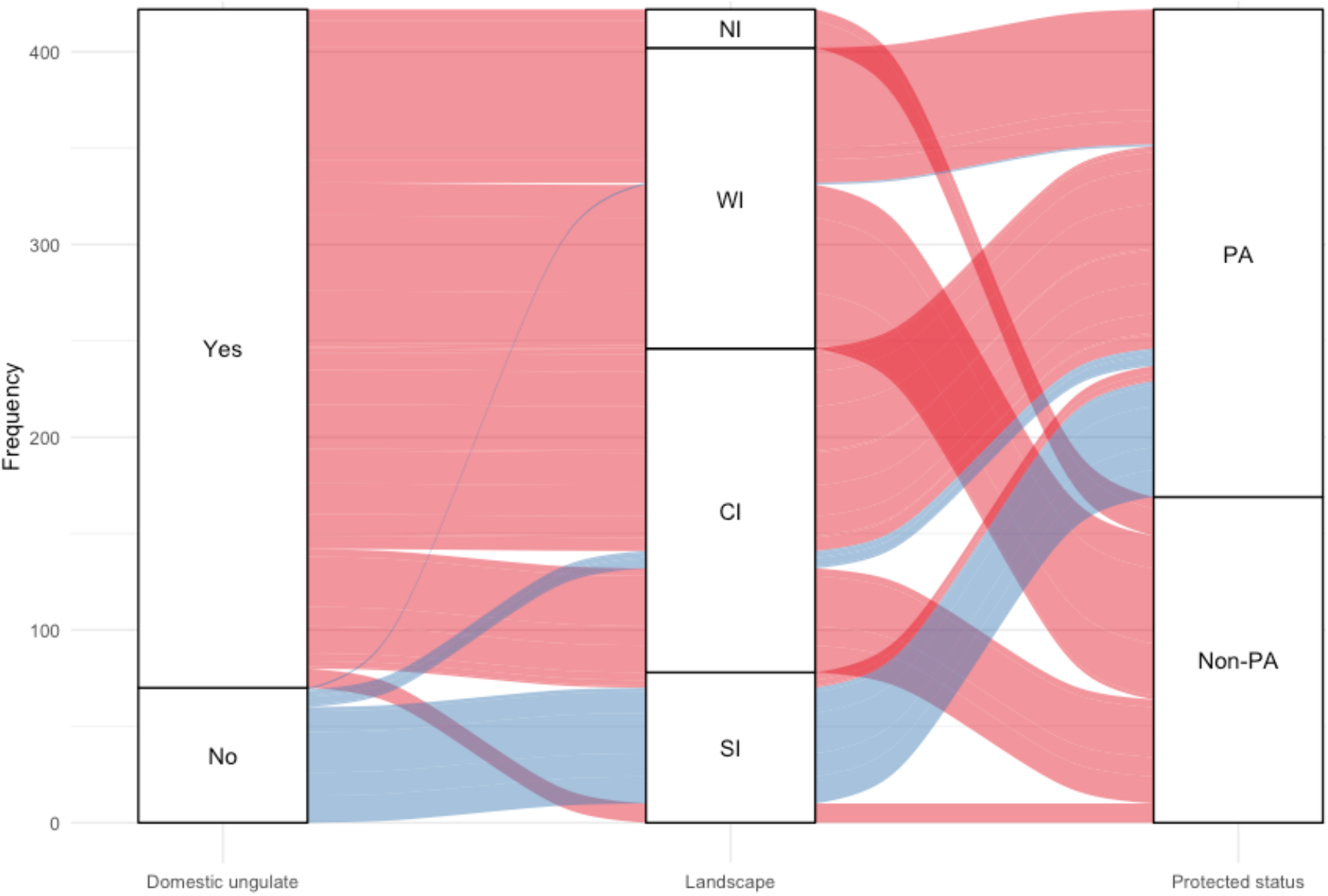
Sankey diagram quantifying the number of links (relative values, *N* = 419) between presence of domesticated ungulates in diet, the landscape (North India, West India, Central India, South India) and protected area status of the site where the faecal sample was collected.

On comparing across landscapes and across protected area status, scaled livestock density within a radius of 100 km of the sites was significantly lower in South Indian sites as compared to sites in the other three landscapes (*F*_*3,25*_*= 7*.*3, p=0*.*0009*, Fig. 5). However, scaled livestock density was not significantly different for sampling sites located inside protected areas compared to those located outside (*F*_*1,25*_*=1*.*6, p=0*.*22*).

**Figure 5.**
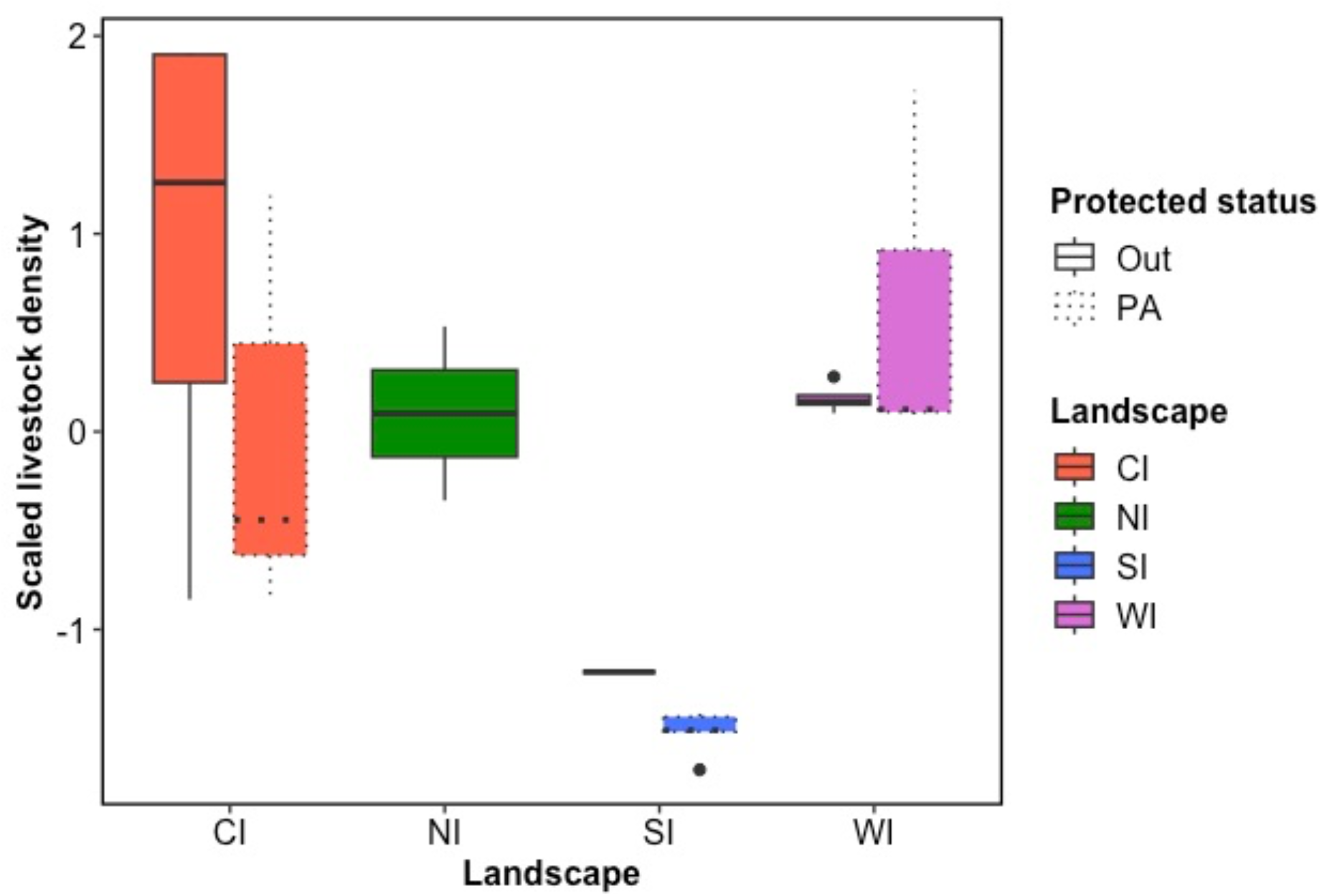
Distribution of scaled livestock density within 100 km^2^ radius of the 30 sampling sites within and outside protected areas across the four landscapes.

## 4. Discussion

We used diet metabarcoding of non-invasively collected faecal samples to describe the diet of *Gyps* vultures from sites located inside and outside protected areas across four landscapes in India. The diet of these vultures was dominated by domesticated large ungulates, except in South India. The observed composition of the diet differed among landscapes and species as well as inside and outside protected areas. This variation was best explained by the differential intake of domestic and wild carcasses. Overall, domesticated ungulates remain the mainstay of *Gyps* vultures diet in most parts of India, the only exception being South India, mainly in response to the very high availability of domestic livestock in the other landscapes. This implies that across most parts of the country, protected areas may not offer a respite from possibility of diclofenac-poisoning and efforts should continue unabated to remove nephrotoxic drugs from veterinary use.

### 4.1 Diet metabarcoding for studying vulture diet

Our study used diet metabarcoding to study variation in vulture diet across a large geographical scale. This was done for two reasons, which made previously used methods based on sampling regurgitated pellets and food remains unreliable. Firstly, *Gyps* species are known to scavenge on nutrient-rich internal organs and soft tissues of large animal carcasses (Houston, 1983; Mundy et al., 1992), and secondly, domesticated ungulate carcasses are often skinned prior to their disposal in many parts of India (Pain et al., 2008). Both these factors make retrieval of undigested hair and bones in the form of regurgitated pellets less likely. This was further confirmed through conversations with experts working in the field, who reported hardly ever seeing pellets containing such remains close to nests/roosts. Our experience in field confirmed the same. Diet metabarcoding overcame these challenges and allowed us to retrieve dietary data from a large proportion (65%) of the faecal droppings collected from varied substrates (soil, leaves, bark, rocks) in differing states of desiccation. We could identify considerable diversity of dietary MOTUs using the approach, while simultaneously identifying the vulture species, which is particularly useful when sampling in areas with multiple species such as roosts and feeding sites. Given the absence of any previous study on vulture diet from the subcontinent, we are unable to assess the sensitivity of DNA metabarcoding relative to previously used methods. However, inability to distinguish between domestic and wild pig, and not being able to use relative read abundance as an index of relative biomass intake of a diet species (see Ghosh-Harihar et al., 2021) represent limitations of the current approach.

### 4.2 Diet composition and spatial variation

The diet of *Gyps* species comprised a wide diversity of mammalian and few avian species, but was dominated by large mammal carcasses (Table A1). Our study represents the first to investigate vulture diet from the subcontinent and for the resident White-rumped and Indian vultures, but a similar preference for large and medium bodied mammals (cow, horses, pig, sheep, goat) has also been documented for Eurasian Griffons in western Europe and Africa (Margalida, 1997; Mundy et al., 1992). We detected several carnivores in the diet, including large cats such as tiger and common leopard, wild dog and jackal, as has been documented from the Great Caucasus where Eurasian Griffons were found to feed on grey wolf, golden jackal and red fox (Karimov and Guliyev, 2017). The presence of a wide diversity and considerable number of dietary MOTUs in the observed diet of the four *Gyps* species also suggests a broader dietary niche breadth for these large-bodied raptors, which are otherwise known to specialize on large ungulate carcasses (Buechley and Şekercioğlu, 2016).

The diet composition of *Gyps* vultures also showed marked spatial variation among landscapes, across sites located inside and outside protected areas as well as between species (Fig. 2, 3). The significant dietary difference between species could be attributed to samples of White-rumped vultures, half of which came from south Indian protected areas with ∼88% showing only wild species in diet. The other species were largely sampled from the other landscapes and had a diet dominated by domesticated ungulates (Fig. 3). The differences between landscapes and across protected areas and outside were best explained by the relative intake of domestic cow and buffalo and chital, the most frequently encountered wild diet species (Fig. 3). The much higher frequency of samples with only wild species in south Indian protected areas (Wayanad and Mudumalai tiger reserves) could not be explained by higher availability of wild prey. Densities of large wild ungulates (chital, sambar, nilgai and gaur), which are the dominant part of vulture diet, are very comparable across well-protected parks sampled in south, central and west Indian landscapes (Jhala et al., 2018, see Table A5). However, availability of domestic ungulate carcasses is considerably lower in south India compared to other landscapes both inside and outside protected areas (Fig. 5), and is understandably the best predictor of domesticated ungulates in diet (Table 1). Furthermore, much higher percentage of rural households consume beef/buffalo meat in the southern Indian states (Kerala-13%, Tamil Nadu-4.8%, Karnataka-2.2%) than the consumption in the other landscapes (Rajasthan (WI) & Madhya Pradesh (CI)-0.3%, Himachal Pradesh (NI)-0.1 %) (NSSO, 2013). This might further reduce the amount and relatively consistent availability of domesticated ungulates carcasses for vultures in southern India.

### 4.3 Livestock and protected areas

The two critically endangered resident *Gyps* vultures are known to range over very large areas based on the few records available (Gilbert et al., 2007; Ram et al., 2022), with the reported home ranges lying between 303 and 68,930 km^2^ for White-rumped vultures and 4,265 to 6,897 km^2^ for Indian vultures. Based on a single study in Gujarat, White-rumped vultures move an average distance of nearly 50 km (max.-91.57 km, min.- 23.6 km) every day while Indian vultures show a mean daily movement distance of 92 km (max.-153 km, min.-32 km) (Ram et al., 2022). The two migratory species, Eurasian Griffon and Himalayan Griffon move over 110 km per day. These data suggest that the relatively smaller protected areas (the largest protected area we sampled is Kanha-941.8 km^2^) in India are inadequate to contain the entire home ranges of these wide-ranging species. This in addition to the presence of livestock grazing within most of these protected areas increases the probability of these species encountering carcasses of domesticated ungulates substantially, as is evident from the diet from most landscapes. Only in southern India, with significantly lower livestock densities in general, did the protected status of a site influenced the presence of domesticated ungulates in the diet. While we do not have data on NSAID prevalence among domesticated ungulates carcasses at the relevant spatial resolution to conclusively establish the risk posed by feeding on domesticated ungulates, the continuing illegal use of banned diclofenac and legal use of other toxic NSAIDs such as aceclofenac and nimesulide is indicative of the risk posed by a diet dominated by domesticated ungulates.

Previous research from parts of India other than the south suggests that the populations of resident *Gyps* vultures from and close to protected National Parks have not only experienced lower declines but continue to show higher abundances at present (Prakash et al., 2019). We sampled most of these populations in central and western India and all show an overwhelming presence of domesticated ungulates in faecal samples (Fig. 4). There could be various reasons behind the buffering effect of protected areas on vulture populations. Firstly, many of these areas present rugged terrain with suitable habitats for cliff-nesting Indian vultures. Furthermore, a significant proportion of the livestock grazing in these protected areas are feral cattle (Jhala et al., 2021; Kolipaka et al., 2017; Rasal et al., 2022; Singh et al., 2011), which do not receive veterinary care and therefore, probably present safe food source for the vultures. Confirming this would require diclofenac prevalence data from carcasses across the landscapes, both inside and outside protected areas. Nevertheless, a recent study suggests that at least two populations of Indian vultures sampled by us (villages in the periphery of Ranthambore National Park and Chambal river gorge near Kota) are experiencing declines relative to populations in Bandhavgarh National Park (McClure et al., 2021). In addition, the citizen science based State of India’s Birds suggests a nationally declining trend for these species, particularly for Indian vultures (SoIB, 2020). Overall, these data suggest that a livestock-dominated diet continues to pose a threat to the remaining vulture populations in the country.

## 5. Conclusions

Our study establishes diet metabarcoding as an efficient method to assess species-specific diet of these obligate scavengers over large geographic scales. The diet of *Gyps* vultures comprised of large mammal species, with domesticated ungulates being present in more than 95% from north, central and west Indian sites. Given the very high availability of domestic livestock in these parts of India, we found that presence of protected areas did not reduce the presence of domesticated ungulates in the vulture diet. In contrast in south India, with much lower livestock density, vultures within protected areas showed a much lower occurrence of domesticated ungulates in their diet than those found outside. These results highlight the importance of continuing the efforts to remove veterinary drugs toxic to vultures from use. This involves testing drugs in use for their effects on vultures, advocacy to ensure such drugs are legally banned, and adequate enforcement and outreach to encourage compliance. Continued monitoring of toxic drugs in carcasses and vulture populations in the wild is critical to ensure these efforts yield the expected outcomes. Based on our results, protected areas in south India appear to pose the least risk of mortality due to toxic veterinary drugs owing to very low incidence of domesticated ungulates in their diet.

## Acknowledgements

We wish to thank the state forest departments of Himachal Pradesh, Rajasthan, Madhya Pradesh, Kerala, Karnataka and Tamil Nadu for providing research permissions for collecting samples. MGH was funded through a NCBS-inStem-Cambridge postdoctoral fellowship. We thank William Amos for advice and supervision of this fellowship. We thank Himanshu Chhattani, Anubhab Khan, Kaushal Patel, Tarsh Thekaekara and Abishek Harihar for help in the field. Meghana Natesh, Ishani Sinha, Harsh Shukla and Abhinav Tyagi are thanked for useful discussions on laboratory protocols and HTS filtering. We thank Anuradha Savand for help with lab work and Abhinav Tyagi for assisting with bioinformatic analysis. Dr Awadhesh Pandit and the Next-Generation-Sequencing facility at NCBS is thanked for processing the samples.

